# Nanotribology of Viruses Reveals Their Adhesion Strength and Modality of Motion on Surfaces

**DOI:** 10.1101/2025.02.03.636328

**Authors:** Charles Ault, Claudia Simon, Irina B. Tsvetkova, Pedro J. de Pablo, Bogdan Dragnea

**Affiliations:** Department of Chemistry, Indiana University, Bloomington, IN 47405, U.S; Boehringer - Ingelheim Pharma GmbH, Germany; Centre International de Formation et de Recherche Avancées en Physique, 077125 Bucharest-Magurele, Romania; Departamento de Física de la Materia Condensada and IFIMAC, Universidad Autónoma de Madrid, 28049 Madrid, Spain

**Keywords:** virus, nanotribology, nanoparticle adsorption, contact mechanics

## Abstract

We studied the friction dynamics of icosahedral viruses adsorbed to solid surfaces to probe their adhesion. Using the lateral torsion of cantilevers in atomic force microscopy to move individual capsids in a liquid environment, we found that the virions tend to roll rather than slide on the surface. In contrast, rigid, ligand-stabilized gold nanoparticles are more likely to combine rolling with sliding under the same conditions. The experiments indicate that the force required to drag the viruses on the surface is four times less than that of AuNPs, while the lateral force work needed to induce virus movement was ∼ 10^4^ kT, ten times less than that of the rigid gold nanoparticles. These results go beyond the paradigm that adhesion of nanoparticles is mainly governed by geometrical factors, such as size and area of contact, highlighting the need to amend modeling approaches to account for mechanically-compliant tribological response of biologically derived nanoparticles.

## Introduction

Infections arising from surface-borne viral pathogens cost an estimated 94 million dollars in treatment and cause half a million deaths in the US annually^1^. The recent SARS-CoV-2 pandemic raised key issues about the adhesion of virus particles to solid surfaces and increased interest in prophylactic approaches to curb virus spreading. In this context, the interaction of biological particles such as viruses with surfaces plays a central role in biohazard mitigation.

The majority of viruses encapsulate their genome inside proteinaceous shells held together by weak non-covalent bonds. At adsorption on a surface, depending on the strength of the adhesive interactions, such soft, mechanically compatible particles partially deform to increase the contact area with the surface with respect to rigid particles^2,3^. There are indications that such malleability may play a role in a nanoparticle’s ability to infiltrate cells and / or cross-cellular synapses^4^. Specifically, adhesive interactions bend the plasma membrane around the particle. In turn, a malleable particle will respond by shape distortion, a phenomenon that has been suggested to modulate the kinetics of cellular uptake^5^.

A physical mechanistic picture of adhesion (and friction) of soft, biologically derived nanoparticles on surfaces, such as viruses and exosomes, is still lacking. There are conceptually straightforward, yet important, questions that remain unanswered. For instance, after landing on a cell’s surface, how much of the virus diffusion on the surface is by rolling vs. sliding until they find an entry point? The answer to this question could inform future kinetic models of receptor mediated endocytosis and the interpretation of single particle tracking experiments in virus entry. Another problem is the stability or metastability of the virus adsorbed on a surface. Its understanding could potentially lead to new antiviral surface chemistries and better separation methods in the preparation of vaccines. However, while adhesion of solid nanoparticles at the air/substrate interface has been studied at microscopic scales^6–10^, the dynamic response of mechanically compliant biological particles to adhesion in a physiological environment has been far less explored^11^. To address this gap, here we have turned to an atomic force microscopy (AFM) lateral force manipulation approach, which we applied to the study of two model viruses adsorbed on dissimilar surfaces.

When two solids are near contact, attractive intermolecular forces cause the bodies to stick together at contact points^12,13^. The force required to overcome these attractive interactions is termed the adhesion force. Adhesion depends on interfacial properties, including, but not limited to, roughness, contact time, deformation, specific interactions, impurities, presence of water, and ambient conditions^12,14^. All of these interactions are also involved in the resistance to sliding motion between two bodies (called friction).

Adhesion and friction (or lubrication) are some of the most familiar, studied, and technologically important macroscopic phenomena. However, how they emerge from microscopic interactions has been a puzzle until recently^15^. In recent decades, advances in instrumentation, particularly the development of AFM, have extended the study of adhesion interactions to nanoscopic systems^16–18^, through direct observation of the role of local properties such as physical defects and strain, and the interrogation of single nanoparticles^19^. These innovations led to an improved molecular-level understanding of adhesion and friction and the development of finer-grained mathematical models^20^.

In surface-adsorbed particles for which constitutive interactions are similar in magnitude to adhesion interactions, the equilibrium shape of an adsorbed particle is the result of balancing elastic and adhesive interactions^21^. The outcome of this energetic tug of war is, generally speaking, the concern of contact mechanics^22,23^. The small shape changes of an adsorbed virus particle with respect to its nominal shape in a homogeneous environment measured by cryo-electron microscopy or X-ray crystallography can be reliably observed *in situ* by topographic AFM mapping^3^.

In recent decades, AFM has been shown to be particularly suitable for interrogating the physical properties of viruses^24,25^ and virus-derived particles^26^. In particular, AFM makes it possible to analyze individual particles in liquid with subnanometer resolution, providing three-dimensional topographic data with real-time capabilities, and facilitating the application of controlled forces^27^. Through the lateral torsion of an AFM cantilever of known geometry and elastic moduli, quantitative forces can be applied to laterally manipulate nanoscopic objects on a solid surface^28,29^ or to collect tribological data^30,31^. For example, the energy required to remove surface-adsorbed functionalized carbon nanorods has been estimated by this method^32^. Moreover, early demonstrations of lateral force manipulations of biomolecular material under ambient conditions included severing surface-adsorbed DNA strands^33^ and stretching of tobacco mosaic virus filaments across a solid substrate^29^. The large aspect ratio of these filamentous molecules benefited from a reduced number of degrees of freedom compared to that of other shapes, which facilitated the interpretation of the AFM torsional data. However, it was not clear until now whether the same approach could be translated to the problem of adsorbed spherical, mechanically compliant nanoparticles in a liquid.

AFM lateral manipulation of rigid spherical particles was demonstrated in air, including silica nanoparticles^34^, polystyrene microspheres^35^, and gold nanoparticles^36^. Mathematical models of the interaction between an idealized AFM probe and rigid micro and nanospheres have been developed^37,38^, most of which are applicable to particles orders of magnitude larger than the most common virus capsids. It is likely that at submicroscopic scales such models would reach their validity limit. In addition, they apply only to rigid particles in an inert gas atmosphere. Their accuracy becomes questionable when the dynamics of surface-adsorbed virus particles are described because of particle compliance and strain under friction and adhesion, and because of the solvent (aqueous solution).

Unlike most similar studies on rigid microparticles, the experiments described here were carried out in an aqueous solution, an environment that drastically alters adhesion. This unexplored setting has practical relevance for virus and extracellular vesicle interactions with surfaces, and could be extended to biological interfaces^39–41^. Thus, this work seeks to provide benchmark results for exploring friction at the nanoscale in an aqueous solution. Specifically, we implement the lateral force manipulation method to carry out the study of static and dynamic aspects of the surface adhesion of two icosahedral virus species, the brome mosaic virus (BMV) and the murine polyoma virus (MPyV), and, for comparison with a solid nanoparticle standard, of Au nanoparticles (NP) adsorbed from liquid onto two different, nearly atomically flat substrates, polar mica and nonpolar, highly-oriented pyrolithic graphite (HOPG).

## Experimental

### Sample Preparation

BMV capsids were prepared according to the agroinfiltration method described in ref.^42^. Yeast–derived MPyV particles were prepared according to the method described in ref.^43^.

Au NPs with diameter 80 nm were synthesized by the polyol method introduced in 1989 by Fi ‘evet, Lagier, and Figlarz^44^, supplemented by an etching step^45^. This approach results in nearly perfect spherical particles a few tens of nanometers in diameter and an overall reduced size heterogeneity compared to other colloidal methods; see supporting information (SI) document, Fig. S2. After purification, NPs are dissolved in pure water and have the surface coated with a thin layer of poly(dimethyldiallylammonium chloride) (PDADMAC) – a hydrophilic, positively charged polymer. Note that since this method yields good results only for particles above ∼ 30 nm diameter, a different synthetic approach is needed for smaller particles.

Au NPs with an average diameter of 24 nm were synthesized according to seed mediated growth in the presence of cetyltrimethylamonium bromide (CTAB)^46^ - hydrophobic surfactant with polar groups. The average particle diameter, size distribution, ligand thickness and dispersion in solution were verified by imaging on a JEOL 1010 TEM and DLS (Fig. S2 in SI).

We note that the difference in the surface ligand chemistry between the 80 nm Au NPs coated with PDADMAC and the 24 nm particles coated with CTAB leads to a very tenuous adhesive bond on HOPG of the 24 nm Au NPs, which highlights the importance of the surface ligand chemistry, even in buffer solutions.

BMV samples were prepared for AFM analysis similar to ref.^3^, by first freshly cleaving a 1 × 1 cm piece of the atomically flat substrate. The substrates were either HOPG or mica. Then, 40 *µ*L of a 0.05 mg/mL purified virus solution at ∼ 10^11^/cm^3^ particles in SAMA buffer (50 mM NaOAc, 8 mM Mg(OAc)_2_, pH 4.5) was deposited onto the substrate and incubated for 30 minutes at room temperature. Following incubation, the samples were placed in the AFM liquid cell for imaging and manipulation. The samples, unlike in the study of bacteria or biomolecules that exhibit strong surface adhesion, were not rinsed because the low concentration of virus particles present in solution at equilibrium does not impede AFM function, and because rinsing will perturb the adsorbed/free equilibrium.

MPyV samples were prepared by first freshly cleaving a 1 cm x 1 cm piece of the substrate, then incubating 40 *µ*L of a solution of 0.05 mg/mL capsid in MPyV storage buffer – 50 mM Tris (Tris(hydroxymethyl)aminomethane) at pH 7.4, 200 mM NaCl, 5 % glycerol, for 30 minutes. The MPyV storage buffer was then blotted and replaced with pH 4.5 SAMA buffer before placing the sample in the AFM liquid cell for imaging and manipulation.

For samples in which BMV and MPyV were co-adhered for direct comparison, the substrate was first freshly cleaved, then MPyV was incubated for 30 minutes as described above, then the sample was dried and 40 *µ* L of BMV solution was immediately added and incubated for 30 minutes.

### Determination of cantilever spring constant

Experiments were performed and data collected using an Asylum Cypher S atomic force microscope running proprietary IgorPro-based (Wavemetrics, Inc) software from Aylum Research (Oxford Instruments, Ltd). The AFM probe was an oxidatively-sharpened BioLever-mini™ BL-AC40TS silicon/silicon nitride cantilever. The cantilever had a nominal size of 38 x 16 x 0.2 *µ*m, a 7 *µ*m tetrahedral probe with tip radius of 8 nm, and nominal resonant frequency of 25 kHz. All data were collected in dynamic mode in solution. The actual spring constant of each cantilever (*K_n_*) was determined using the Sader method^47^. Typically, the normal spring constant was found to be ∼ 0.09 ± 0.03 N/m, in agreement with the manufacturer’s specifications.

The lateral spring constant (*K_l_*) was determined using a standard method^32^ based on geometric conversion^48^ as shown in equation 1.

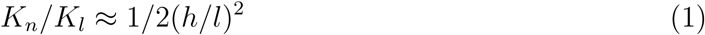

where *h* and *l* are the height of the probe and the length of the cantilever, respectively. It should be noted that formula (1) provides a quick but rough estimate of the lateral force constant. This approximate method has been used previously^32,49–51^. However, different approaches exist, which would be preferable when higher accuracy is desired^52,53^.

The sensitivity of the photodiode to lateral torsion of the cantilever was calibrated by scanning the 100 nm height step of a commercially available milled grid. The abrupt increases in lateral force as the probe crossed the step was recorded. The slopes of lateral deflection vs horizontal distance were used to calibrate lateral force sensitivity and convert the measured lateral signal voltage to torsional displacement and then, using the lateral spring constant, into lateral force (see SI, Fig. S6 for an example).

### Lateral force manipulation

To perform the manipulations, a topographic image was first collected of the particles adhered to a substrate. Then a particle which exhibited nominal physical features was isolated from nearby particles and selected as the object of investigation. Using the ‘MicroAngelo’ tool in the AFM software, a typical probe path approximately 200 nm in length was prescribed which passed through the center of the capsid of interest normal to the length of the cantilever. A pre-scan along this path was performed in tapping mode during which a topographic profile was collected. From these data, a new probe path was created that passed through the capsid, with the tip starting at a small controllable distance below the particle height and above the surface of the substrate. A manipulation was then performed in dynamic mode with the amplitude feedback loop turned off. As the probe followed this path, the amplitude, normal deflection, and torsional deflection were collected as a function of the probe position. Following manipulation, a new topographic image was collected. Comparison of the first and second topographic images provided the data required to quantify changes in capsid position and/or structure.

In some lateral manipulation experiments, to facilitate direct comparison, BMV and MPyV particles were co-adhered alongside one another on the same substrate to create a heterogeneous particle sample. This practice ensured identical environments and probing for both species.

### Data Processing

Topographical data was processed using WSxM 5.0 Develop 10.2^54^. For each manipulation, the topographic image generated by the z-piezo readout was investigated prior to and following the manipulation. Both files were first cleaned using the multi-flatten filter tool with the ‘line’ subtraction type and substrate area parameter set at 60-80 %. The corrected images were then aligned using the *align images* tool. The topography of the manipulated particle was taken along the axis of motion and the traces were overlayed to determine the distance of particle displacement. All other data processing was performed with OriginPro 2023. The pre-scan topography, deflection, and amplitude were plotted vs. path length without any processing. The lateral force was determined by converting the photodiode signal into the torsion distance at the tip as seen SI, Fig. S6, and then multiplying this distance by the lateral spring constant. The normal force could also be determined by multiplying the deflection by the normal spring constant. The lateral force trace vs. distance was integrated using the ‘integration’ gadget of OriginPro to determine the lateral force work.

## Results and Discussion

The first step in an AFM lateral force manipulation experiment is the identification of adsorbed particles in dynamic (AC) mode. Figure 1(a) shows an example of a spread of MPyV particles on mica, in solution. The geometric building blocks that give the capsid its knobby aspect are called capsomers.

**Figure 1:**
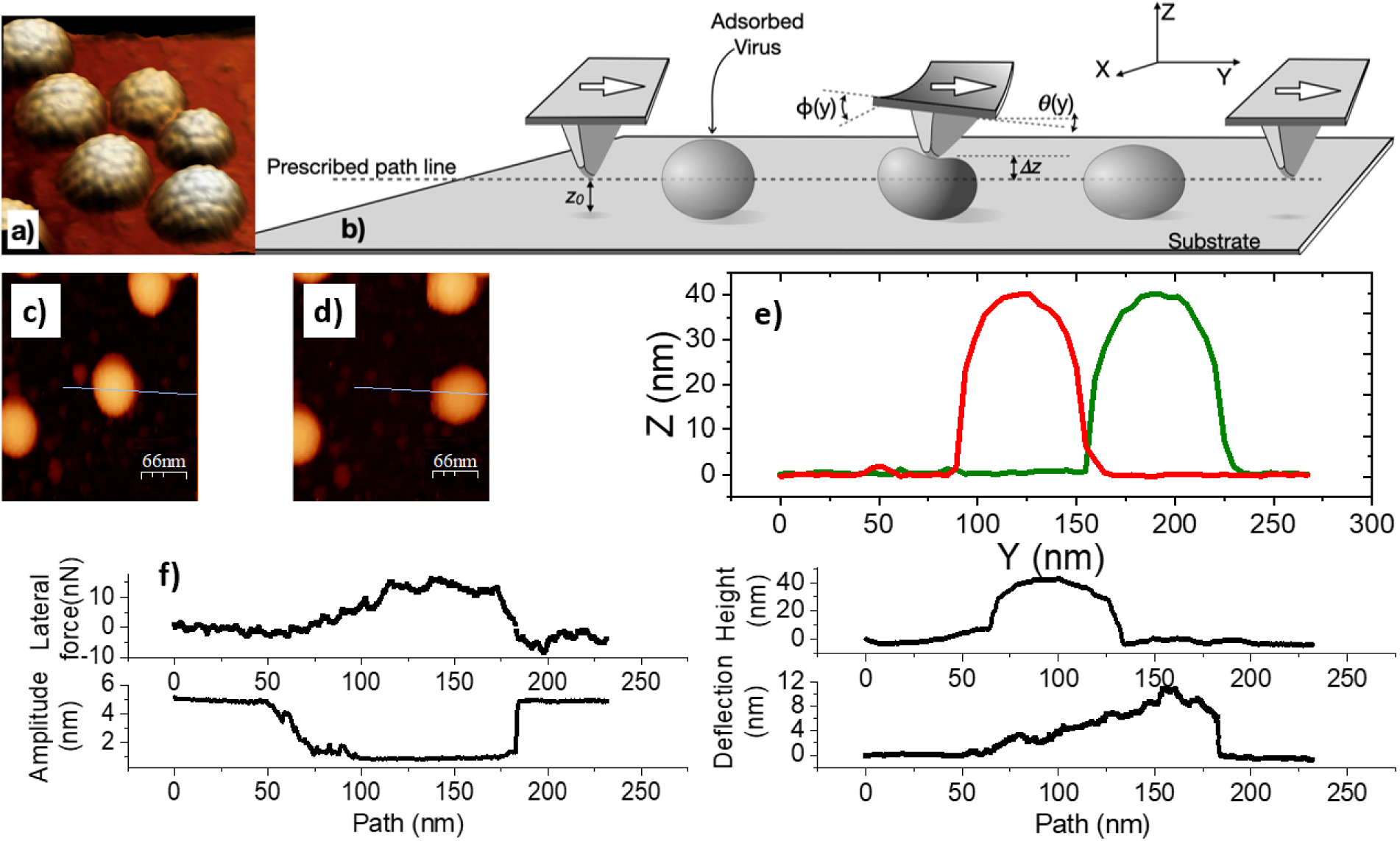
Principles of lateral force manipulation by AFM, illustrated with typical data for MPyV particles, adsorbed on HOPG, in buffer. (a) AFM topography of a group of MPyV capsids adhered to HOPG. (b) Cartoon of the lateral force manipulation experiment. *ϕ* is the torsion angle, while *θ* is the deflection angle. Δ*z* is the linear deflection associated with *θ*. *z*_0_ is the tip position above the surface, on the prescribed path parallel to the substrate. (c) and (d) images before and after manipulation of a virus particle by the tip moving along the blue line. (e) Topographic plot profiles of the particle before (green) and after (red) manipulation. (f) Lateral force, oscillation amplitude, topography, and deflection profiles for a particle undergoing displacement under lateral force manipulation.

Members of the polyoma family are distinct from other icosahedral virus structures obeying the Caspar-Klug quasi-equivalent geometry^55^ by having all capsomers internally organized as pentameric oligomers of the protein VP1 (Fig. S1). In contrast, the BMV capsid has a T=3 canonical Caspar-Klug structure^55^ formed of capsomers that are internally organized as hexameric or pentameric oligomers of the same coat protein (Fig. S1).

After imaging and particle selection, a linear path for tip motion was defined (dashed line, Fig. 1(b)) which crossed the adsorbed particle of choice. The paths run perpendicular to the longest axis of the cantilever, through the center of the particle (blue line in Fig. 1(c)-(d).) At the start, the tip was placed at a small vertical distance (*z*_0_ ∼4 nm in this case) from the sample surface, Fig. 1(b). The feedback loop for the probe actuator that drives the motion normal to the substrate was disabled during the tip motion along the manipulation path. Tip velocities were typically adjusted to ∼ 100 nm/s; slower values gave raise to artifacts from thermal drift, higher values frequently dislodge and desorb the particle. From pre-scan and post-scan topographical images (Fig. 1(c)-(d)) changes in particle position and height can be evaluated (Fig. 1(e)).

Throughout the lateral manipulation experiment, the tip oscillates vertically (∼ 5 nm) at resonance. Changes in the amplitude of tip oscillation inform on the nature of the contact between the sample and the tip (Fig 1(f), bottom left). Specifically, when the AFM tip is in full contact with the particle, the oscillation amplitude is reduced to zero and we expect it to have little influence on manipulation.. In fact, the oscillation amplitude accounts for just ∼ 10% of the smallest virus (BMV) size. In this case, this amplitude would apply a maximum force of *F* = 300 pN to the virions, inducing a deformation of *F/k_v_* = 1.5 nm, where *k_v_*is the stiffness of BMV^56^. This deformation represents only ∼ 5% of the size of the virus. However, if the particle and tip are not in close contact, this amplitude reduction is only partial. Thus, the tip amplitude is a qualitative descriptor of the tip-particle contact tightness and, indirectly, an indicator of possible particle deviation from the prescribed path (like when the particle spins away from the probe). Parameters recorded simultaneously along the path included the oscillation amplitude, the deflection of the cantilever, and the torsion angles (Fig. 1(f)). Together, this set of variables provides a set of clues about the nature of motion, as discussed below.

To examine a specific example as a preview, let us consider the data obtained from a MPyV particle in liquid on HOPG (Fig. 1). First of all, we found that the particle can be pushed along without dissociating it from the surface into the liquid. This is important to note because it was not obvious before these experiments that the particles would not detach from the surface once they started to move. Particle release sometimes occurs depending on the buffer, substrate, and mechanical experimental parameters, but it is possible to tune those conditions to make dissociation a less likely event. Second, the height of the MPyV particle did not change after manipulation, suggesting that the process generally leaves the particles intact. Third, the distance moved from the initial location is limited to about half of the virus perimeter. After covering that distance, the probe loses contact with the virus particle in the cantilever, regaining the initial torsion and deflection values.

After the deflection and torsion angles *ϕ* and *θ* (Fig. 1(b)) were converted to nm and nN values, respectively (Fig. 1(f)) (see the Methods section), we found that the maximum cantilever deflection (12 nm) is significantly lower than the nominal virus height (46 nm). Since the probe deflection takes positive values throughout probe/particle contact, we deduce that the tip climbs on the particle during manipulation. However, if MPyV behaved like a rigid particle, the estimated deflection would have been ≈ *R*(1 +cos *α*) − *z*_0_ where *z*_0_ ≈ 5 nm is the vertical offset from the surface, *R* ≈ 23 nm and *α* ≈ 0.1 rad. The expected deflection if MPyV were rigid and the tip climbed on top would be ≈ 38 nm, i.e. about 3 times the observed value. Since probe deflection was only 30% of the particle height we infer that, either the tip did not climb all the way over the particle and descended on the other side, in which case we have to assume that the particle was sliding in front of the tip, or the tip did climb but the particle deformed under the tip with a force of ∼ 0.08 N*/*m × 12 nm = 1 nN.

The first hypothesis falls short in explaining the path length characteristics, as we shall see in more detail later. If the virus particle is compressed and the deformation is detectable, do we see something different when rigid particles are investigated under the same conditions? A representative data set for the manipulation of an Au nanoparticle (NP) displaced by lateral force manipulation is presented in Fig. 2.

**Figure 2:**
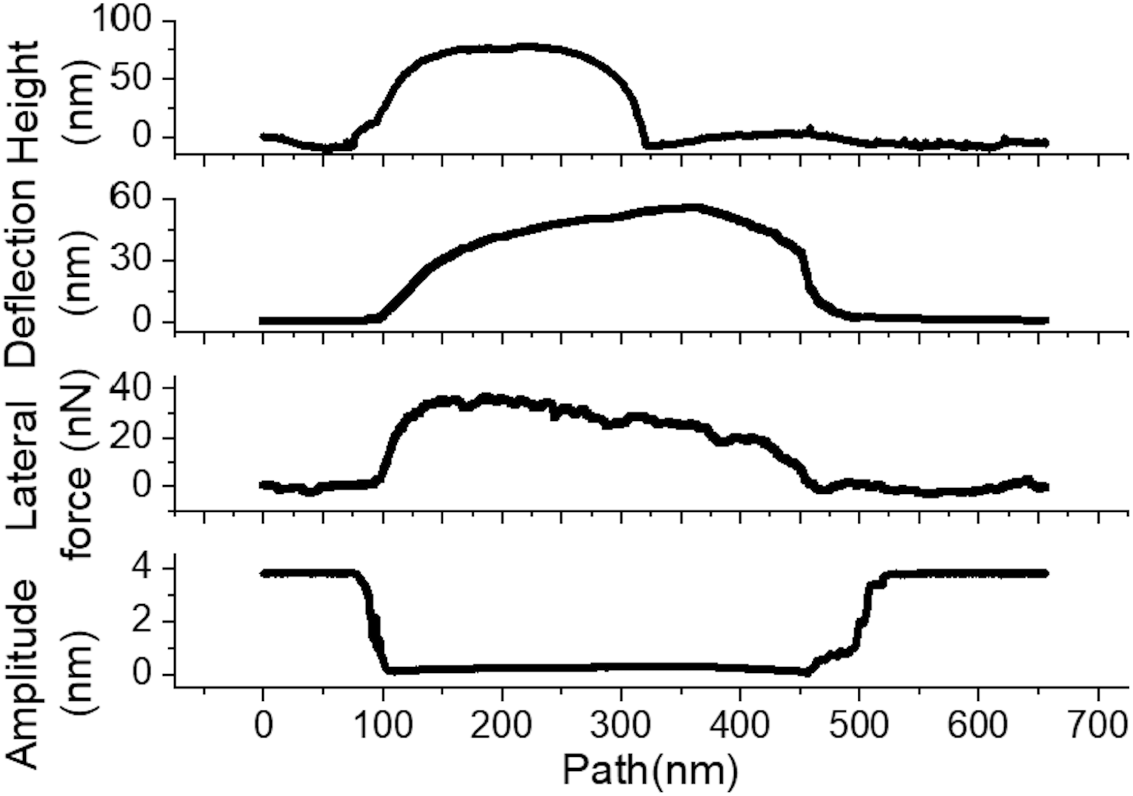
Representative pre- and post-manipulation topography (left) and pre-scan profile, deflection, lateral, and amplitude data (right) for the manipulation of an 80 ± 5 nm Au NP on HOPG.

Indeed, in this case, the maximum deflection reaches ∼ 75% of the nominal particle height which significantly exceeds that obtained from MPyV. This means that the AFM tip climbs almost all the way up the Au NP and that a miscalibration of deflection would not explain the reduced deflection readings on viruses. The most likely explanation is that significant vertical compression occurs during virus manipulation. Two other telling differences between Au NPs and viruses should be noted (Fig, 2): i) The lateral force during the Au NP manipulation was 4× higher than the maximum lateral force on MPyV; ii) the displaced distance (∼ 500nm) was much longer for Au NP, than for MPyV.

To test if these qualitative differences exist when other non-enveloped icosahedral viruses are considered, we have examined the behavior of BMV adsorbed on HOPG, in buffer solution, too. BMV is a single-stranded, positive-sense RNA plant virus that has been studied for nearly 70 years as a model for the broad class of single-stranded RNA icosahedral alphaviruses^57^. BMV particles are about half the size of MPyV, having a diameter of ∼28 nm. Its protein coat is made up of 180 proteins, copies of the same gene (SI, Fig. S1). Due to a mosaic of charged polar and non-polar aminoacid residues at its surface, BMV readily adsorbs from buffer onto a variety of substrates, and is relatively straightforward to image by AFM at spatial resolutions better than 5 nm^3^.

Similarly to MPyV, BMV was displaced along the surface without desorption into the liquid. Fig. 3 shows representative lateral force manipulation data for a BMV particle adsorbed in HOPG. After contact with the probe, the lateral force increased to ∼ 8 nN, plateaued, and gradually decreased back to zero. This plateau represents a stationary state in which resistance to movement is balanced by lateral pushing force. The approximate path length under probe contact was approximately half of the virus perimeter, similar to MPyV.

**Figure 3:**
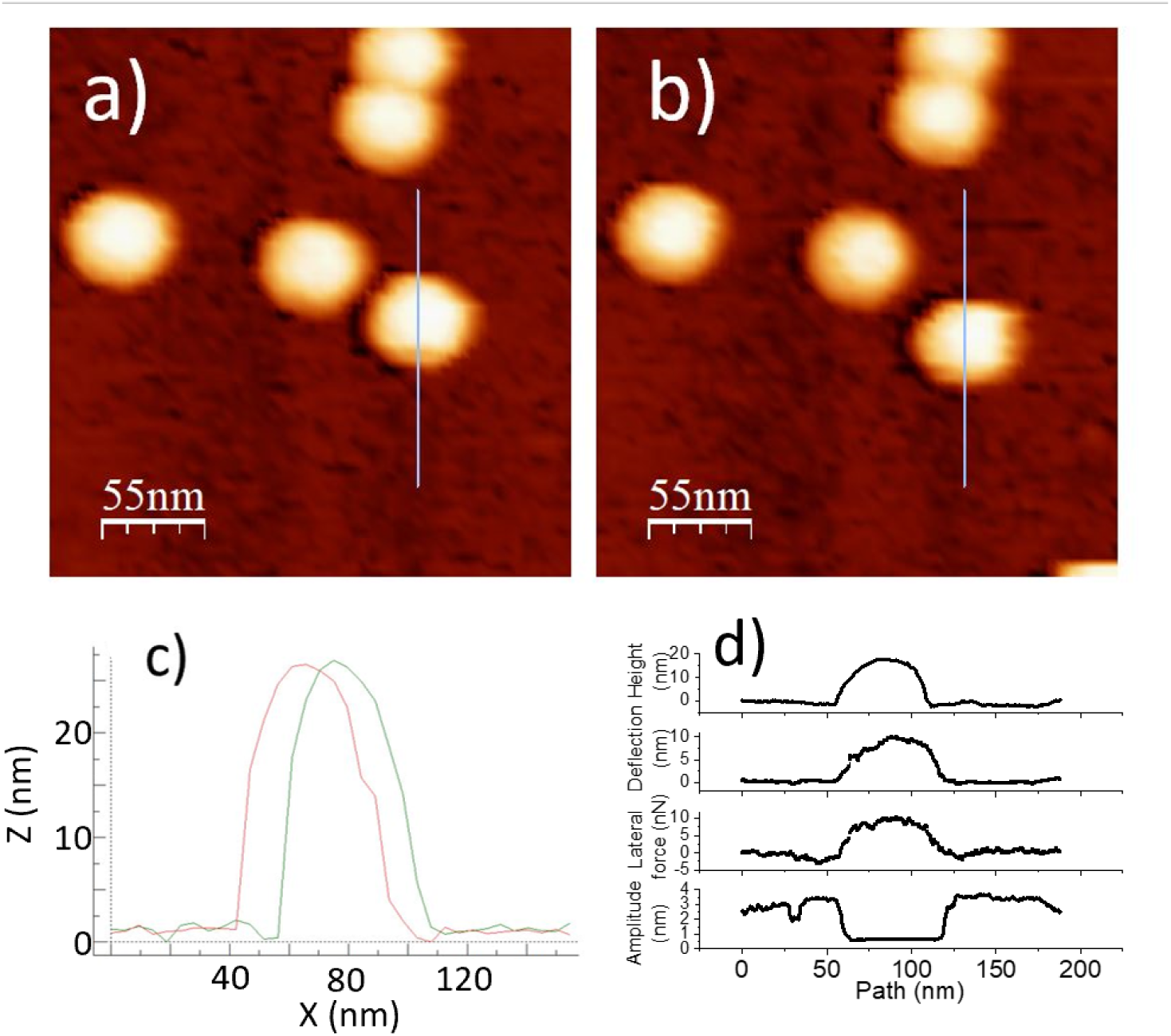
A BMV particle on HOPG to be displaced according to a prescribed probe path during a lateral force manipulation measurement: (a) Before manipulation. (b) After manipulation. (c) Overlay of line profiles, before and after. (d) Topography, deflection, lateral force, and amplitude of manipulation.

As a starting point for further analysis, we refer to previous studies of lateral force manipulation of *rigid microparticles* for which three types of particle motion have been identified: sliding, spinning, and rolling^38,58^ (Fig. 4). It is reasonable to assume that the same type of motion may occur at nanoscale as well. To those types of motion, we add the possibility that a particle be stuck on the substrate while the tip slides on its top surface, while the cantilever deflects up and the particle compresses down. The latter instance has not been reported on microparticles.

**Figure 4:**
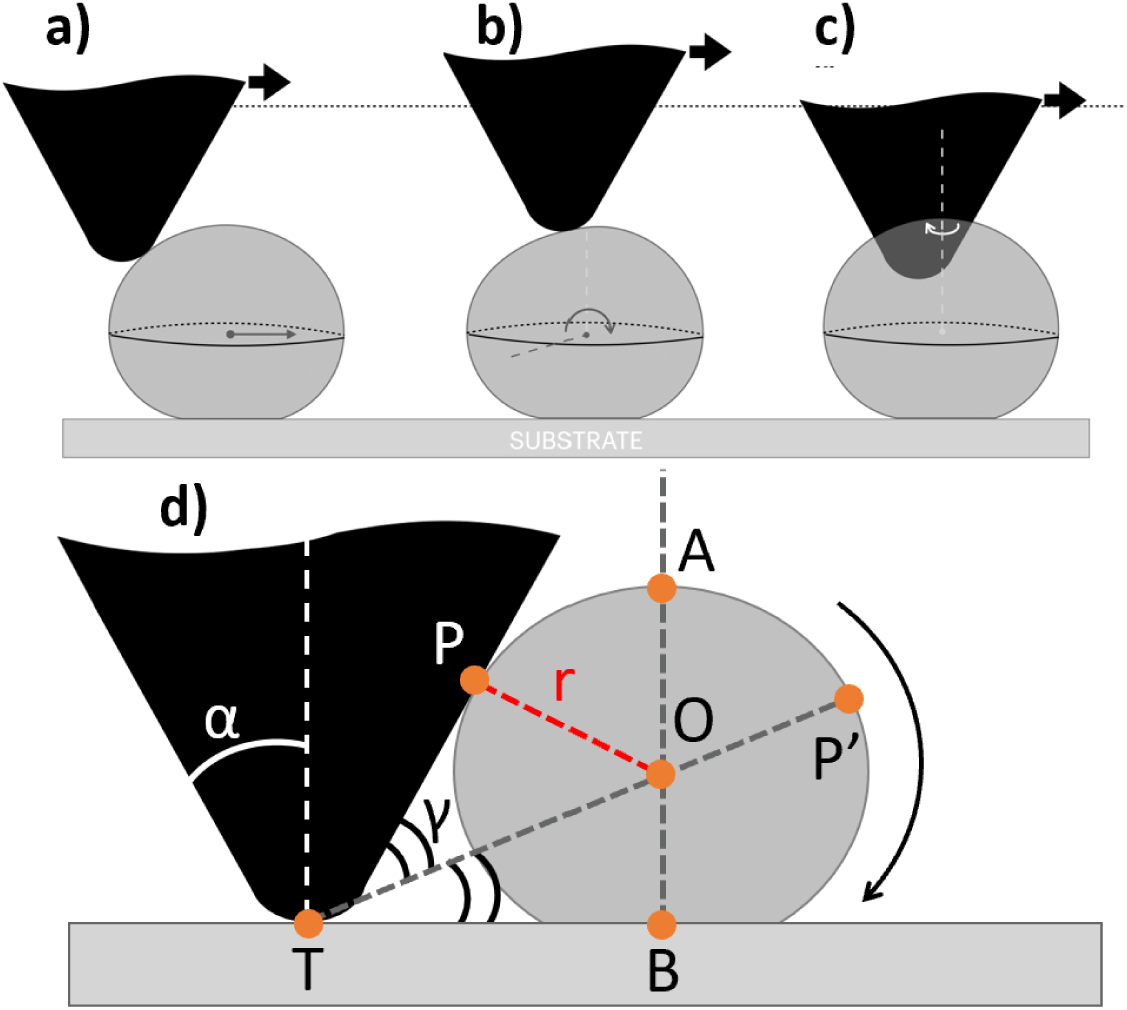
Three motion types for particle undergoing lateral force manipulation: (a) sliding, (b) rolling, and (c) spinning, and diagram of the geometric rolling model (d).

Having access to multi-dimensional data as presented in Fig. 1(f) can help to discriminate between different scenarios of lateral manipulation. This distinction is important to make because each situation informs of different tribological characteristics (e.g. rolling vs. sliding friction, etc.).

We start with particles that spin around the probe, Fig 4(c), because these particles were removed from further analysis. These particles were identified as having a brief lateral and deflection signal during manipulation, the final particle location deviating from the intended probe path, Fig. 5, (unlike in Fig. 3(a-b), where the displacement occurred along the probe path). This situation was usually accompanied by an erratic oscillation amplitude trace, Fig. 6. Together, these results are consistent with the particle spinning around the probe. We note that such instances were also generally associated with lateral force work values substantially smaller than when probe-particle contact was constant and tight, and the direction of particle motion closely aligned with the direction of pushing – see similar observation on microparticles, in Sitti et. al^38^. Thus we have left out of the analysis the particles exhibiting any of the tell-tale signs of spinning around the probe.

**Figure 5:**
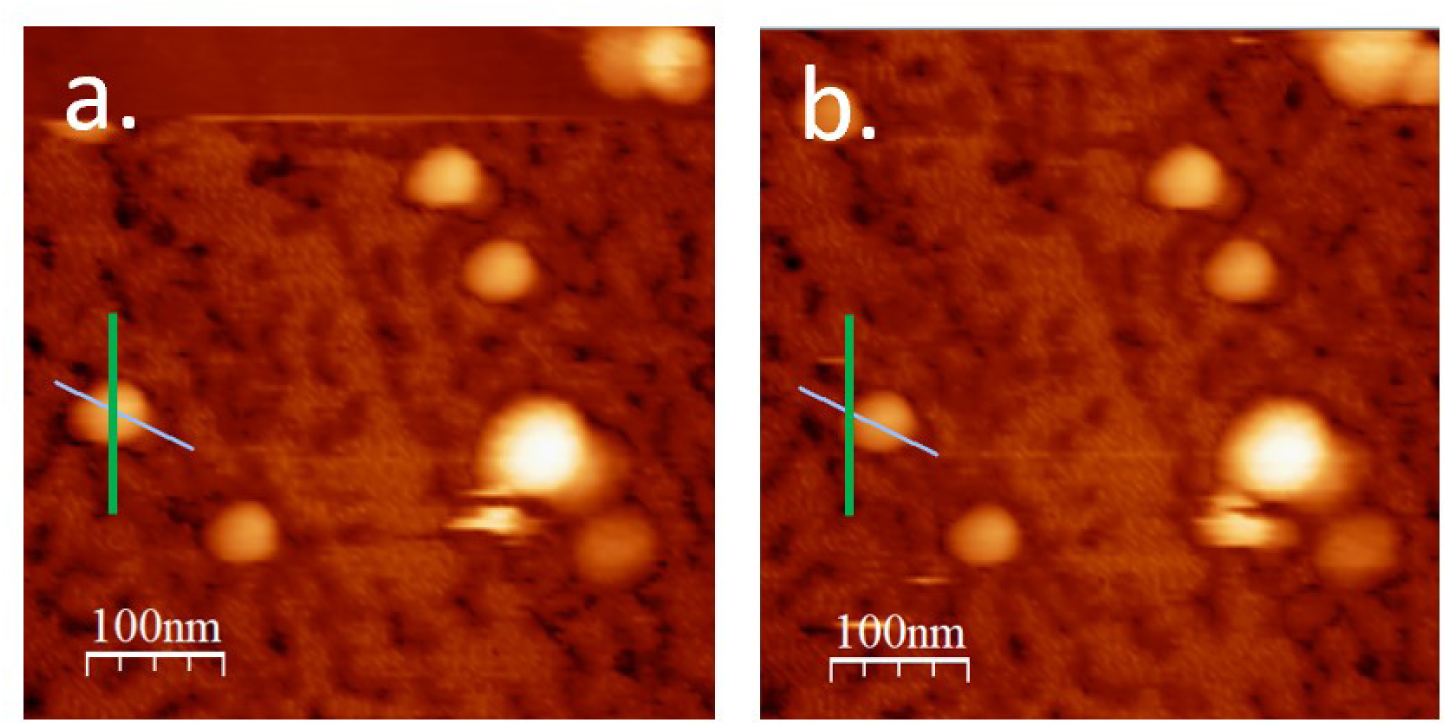
Pre-manipulation (a) and post-manipulation (b) topography of a BMV particle which has undergone spinning during manipulation. Green lines indicate probe path, cyan lines indicate the apparent direction of particle displacement.

**Figure 6:**
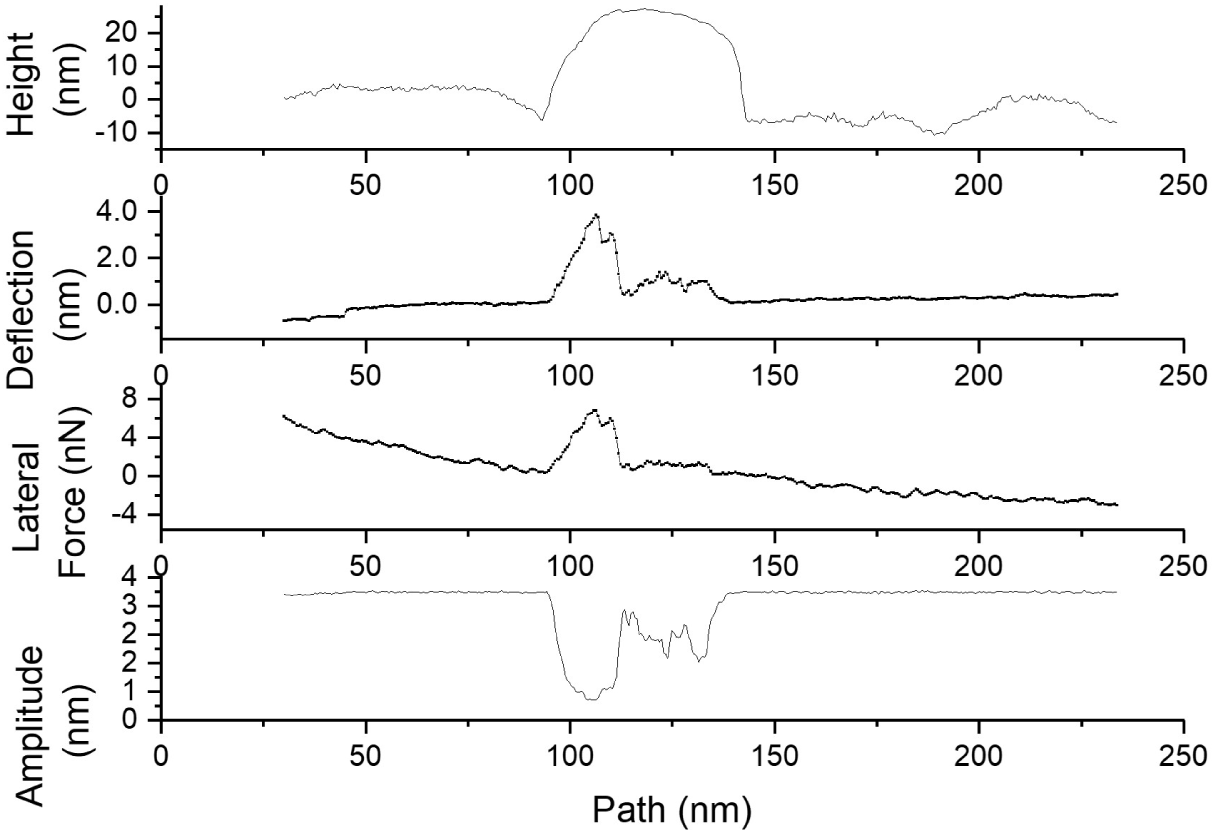
Initial topographic cross-section, deflection, lateral force, and amplitude traces showing characteristics of a particle that exhibits spinning around the probe during manipulation.

Another straightforward scenario to identify is that of a surface-stuck particle. Figure 7 presents data from a BMV particle that has not changed its location on the surface during attempted manipulation. Clearly the particle has not moved on the surface as suggested by neighboring surface features that serve as fiducial markers. In this case, the particle and probe had good contact, but the friction between the tip and particle must have been less than the adhesion between the particle and substrate. As a result, the tip slid across the particle rather than the particle being displaced across the substrate. As we shall see in the following, the number of “stuck” particles can be controlled via the distance of the tip to the surface. Stuck particles could be useful analytically because they can provide independent information on the friction between the probe and the virus.

**Figure 7:**
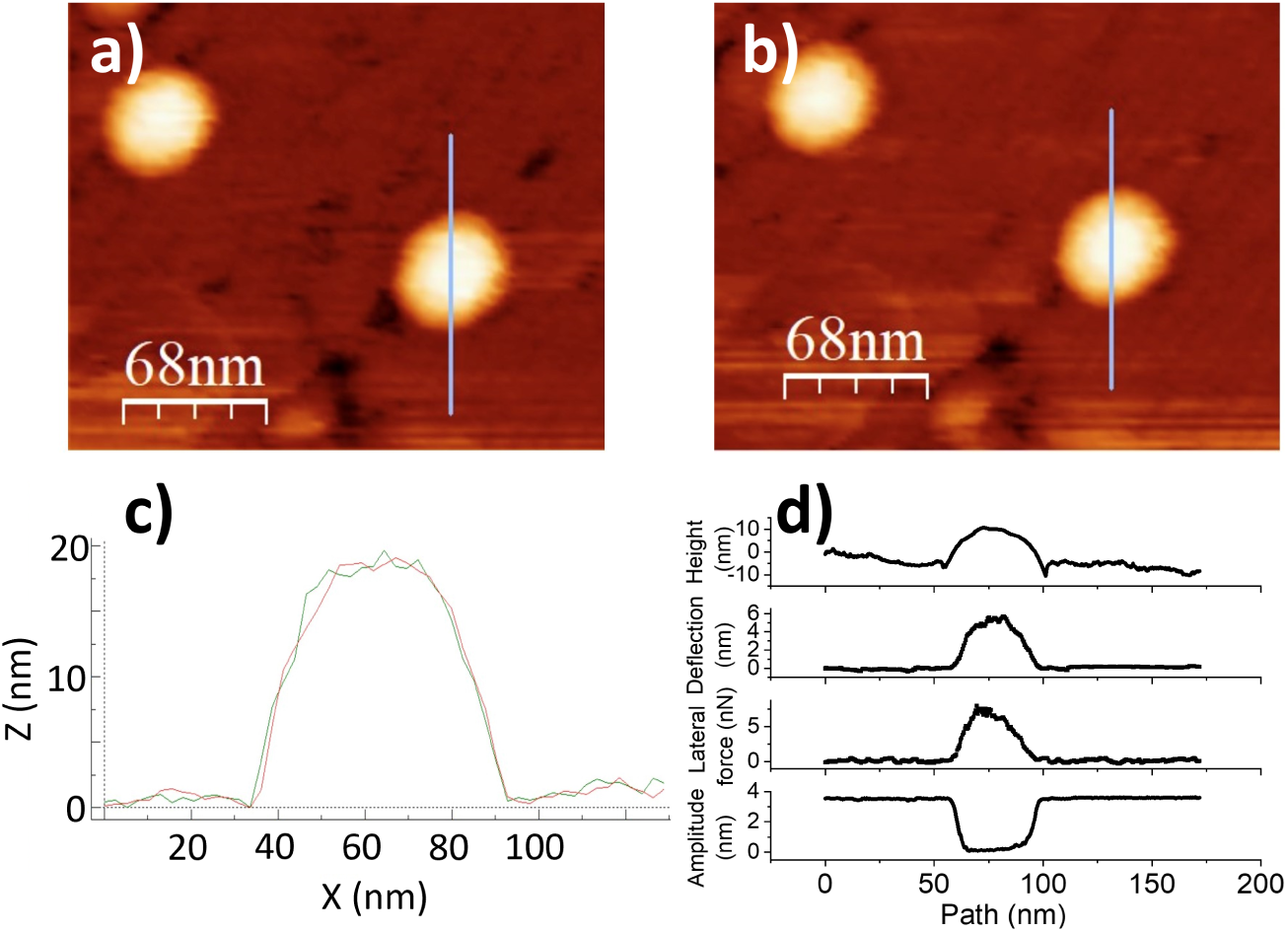
Stuck particle data. (a)-(b)Topographic images of BMV particles adsorbed on HOPG, before and after lateral force manipulation. The blue line indicates the prescribed probe path. In this case, there was no particle displacement by the tip. (c) Single line scan across the particle showing identical topographic profile before and after. (d) pre-scan topography, deflection, lateral force, and amplitude of oscillation at every point along the manipulation path.

We now turn our attention to the remaining two possible mechanisms of motion: sliding (Fig. 4(a)) and rolling (Fig. 4(b)). In the case of sliding, the path length under probe contact can, in principle, be unlimited. Nevertheless, inhomogeneity of surface interactions can kick the particle out of the probe’s way. However, in the case of rolling, there should be an upper limit for the path length, related to the particle diameter. The geometrical model below provides this limit under the following assumptions (see Fig. 4(d)):

- The particle is rolling onto the substrate while the tip is riding on it it without sliding.
- The particle is spherical.
- The probe interacts with the particle via a flat side. The tip, the contact point, and the center of the particle define a plane that contains the assigned linear trajectory and the normal to the substrate through the center of the particle.
- The initial tip position is very close to the surface.
- Particle deformation is negligible.
- There is no sliding between the tip and the particle and the particle and the surface. In other words, adhesives bonds at these interfaces are tenuous with respect to dynamic friction.

The maximum rolling distance (*m.r.d.*) predicted by this model is:

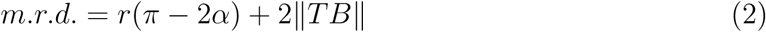

where *α* is the half-angle of the tip pyramid, taken here to be *π/*6 radians.

For ∥*TB*∥ we have:

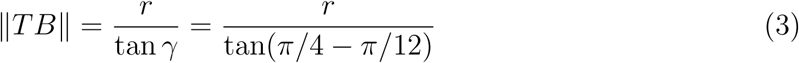

where *r* is the particle radius, and *γ* stands for the *P*^-^*T O* angle in Fig. 4(d)). We obtain:

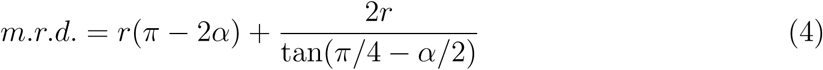

Assuming *r* ≈ 15 nm (for BMV) and *α* ≈ *π/*6, we obtain: *m.r.d* ≈ 84 nm, which is somewhat larger, but close to the experimental value of ∼ 65 nm (Fig 8). Note that a shortening of the effective particle radius may occur due to deformation during manipulation, which is not included in the model. The top limiting value provided by the rolling model clearly sits on the boundary of traveled distances for both BMV and MPyV, Fig 8. In particular, one in 53 BMV particles and none of the 52 MPyV particles have been observed to exceed the *m.r.d.* estimate. However, Au particles frequently exceeded their *m.r.d.* estimate, indicating that a substantial fraction of them presumably moved via sliding.

**Figure 8:**
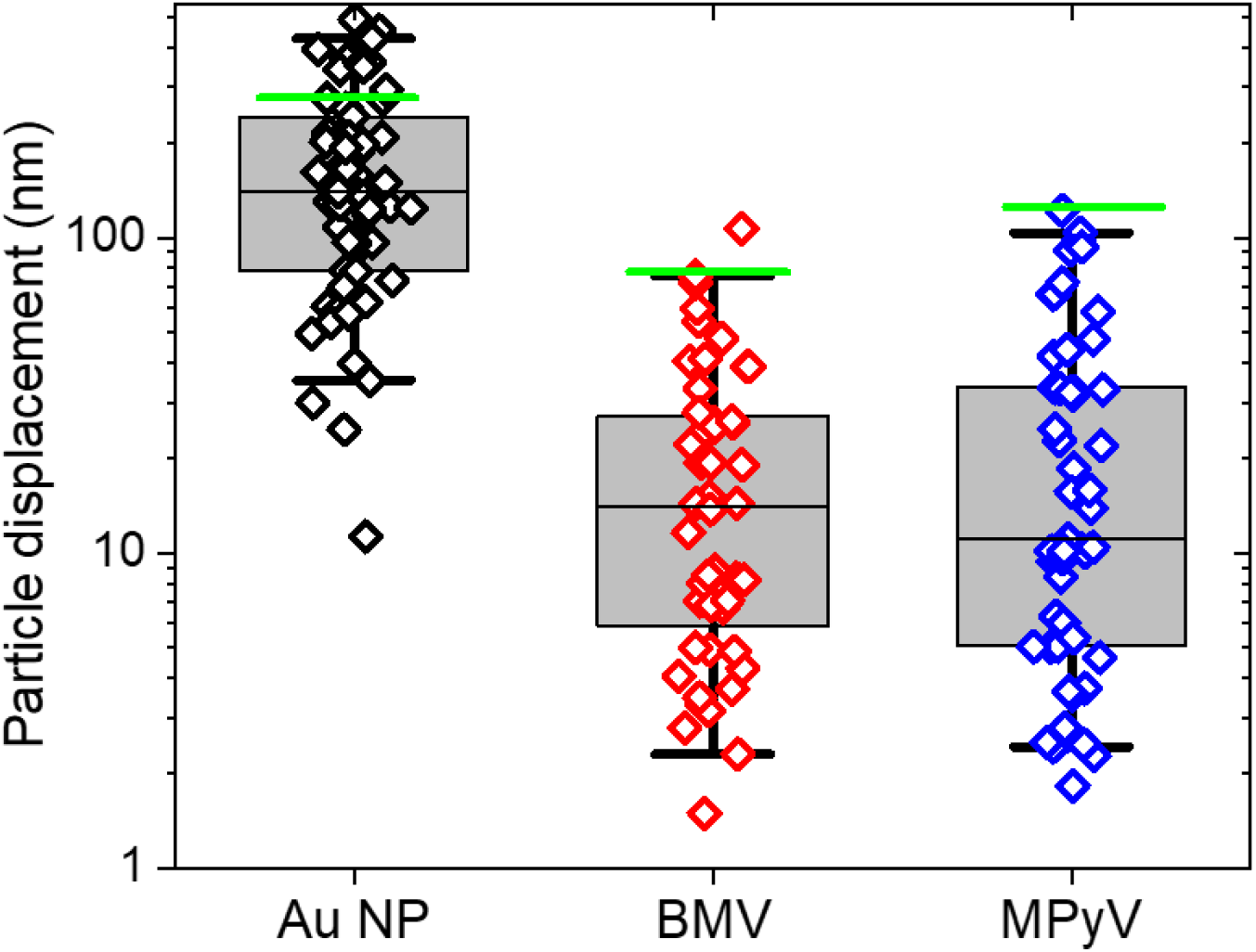
Experimental displacements and calculated maximum rolling distance (green line) for Au nanoparticles (N=62), BMV (N=53) and MPyV (N=52) adhered to HOPG during manipulation in which the probe rides over the rolling particle. Whiskers indicate the middle 95 percent the population, boxes indicate one standard deviation from median of data.

It has already been mentioned that not all adsorbed BMV particles will undergo surface-bound displacement, being “stuck”. It is straightforward to distinguish between displaced particles (Fig. 1(e)) and stuck particles by comparing initial and final locations in the context of surface fiducial markers (Fig. 7). The maximum tip deflection for the stuck particle was ∼ 6 nm, corresponding to a normal force of ∼ 0.6 nN if we consider a stiffness of the cantilever of 0.07 N/m. Under this normal force, BMV is expected to compress ∼ 5 nm^59^, which is significant relative to the virus radius (14 nm). The measured initial height of the stuck particle in Fig. 7 was about 70% of the nominal diameter of a BMV virion.

This low height suggests that the stuck particle might have been structurally altered, most likely at the contact area with the substrate. It is reasonable to expect an increased particle-substrate interaction arising from this alteration, which explains why the particle resisted displacement although the experimental parameters were, in principle, the same as for the displaced BMV particle in Fig. 1(e).

The presence of stuck particles provides an opportunity to compare friction at the probe/particle interface with the rolling resistance. Control over the fraction of mobile particles can be obtained by following the idea that when the tip radius is comparable to the virus radius, the loading angle will vary considerably as a function of the height of the tip as measured from the substrate (*z*_0_ on Fig. 1(b)). Thus, when the tip moves at a distance from the surface that is close to the virus apex height, the tip will merely skim the surface of the virus, exerting reduced forces with respect to the head-on collision case when the tip-substrate distance would be, say, only 5 nm. It follows that there should be a crossover between the tip sliding over the particle *vs.* the tip causing the particle to roll over the substrate as the tip/surface distance decreases. To test this expectation, we have carried out experiments on BMV/HOPG at 5, 10, 15, and 20 nm trajectory depths, Fig.9, where “depth” is understood here as the difference between the height of the particle and the height of the trajectory over the support surface *i.e.d* − *z*_0_.

**Figure 9:**
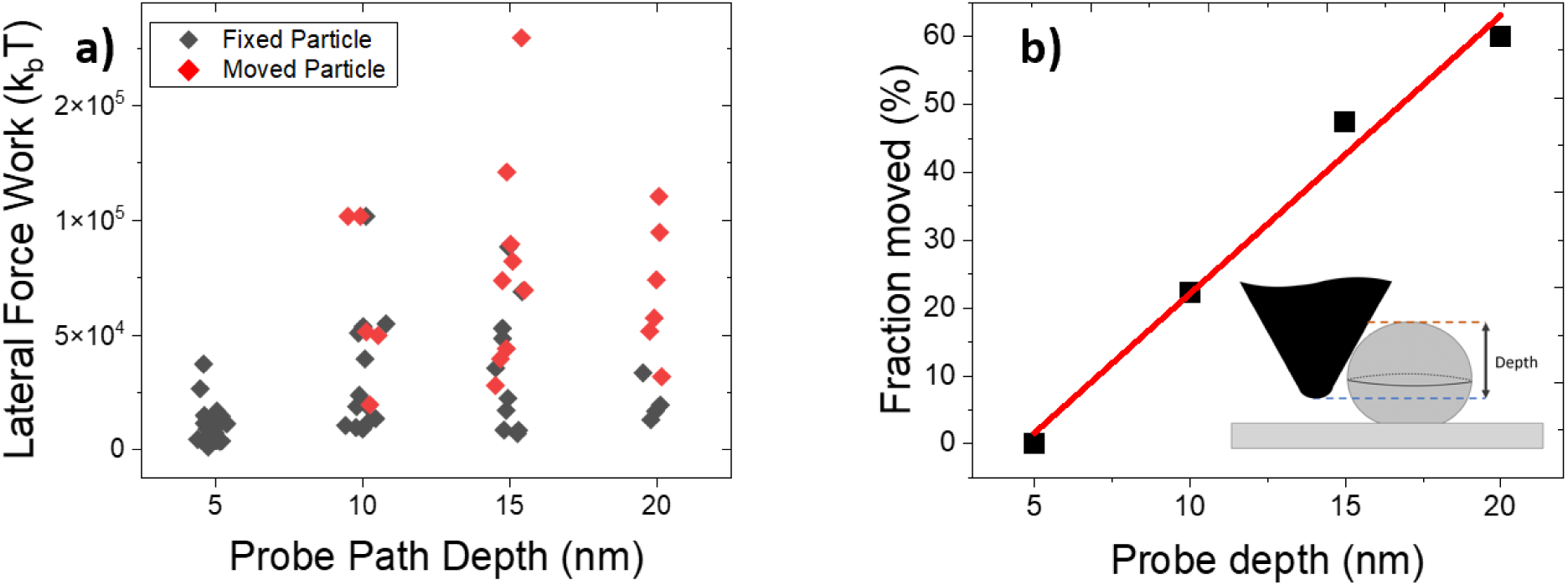
a) Scatter plots of lateral force work for displaced and fixed particles. Boxes and whiskers indicate middle 50 percentile and 90 percentiles of the populations respectively. (b) Percentage of displaced particles as a function of the tip trajectory depth, defined as particle height minus *z*_0_.

For a fixed tip depth, the work of the lateral force is less when the particle is stuck than when it is moving Fig.9(a). As expected from a rolling scenario, displacement occurred more frequently at greater tip depths, Fig. 9(b). Notably, the average lateral force mechanical work for a rolling particle was about 7.3 × 10^4^ kT. This dissipated energy is an order of magnitude higher than the estimated free energy of assembly^60,61^; presumably, most of it is consumed by breaking surface-particle binding, without measurably perturbing particle structure. This observation indicates a large activation barrier to disassembly by mechanical/adhesive forces. However, this barrier is reachable as suggested by repeated manipulation of a BMV particle, which led to an obvious loss of material (Fig. 10).

**Figure 10:**
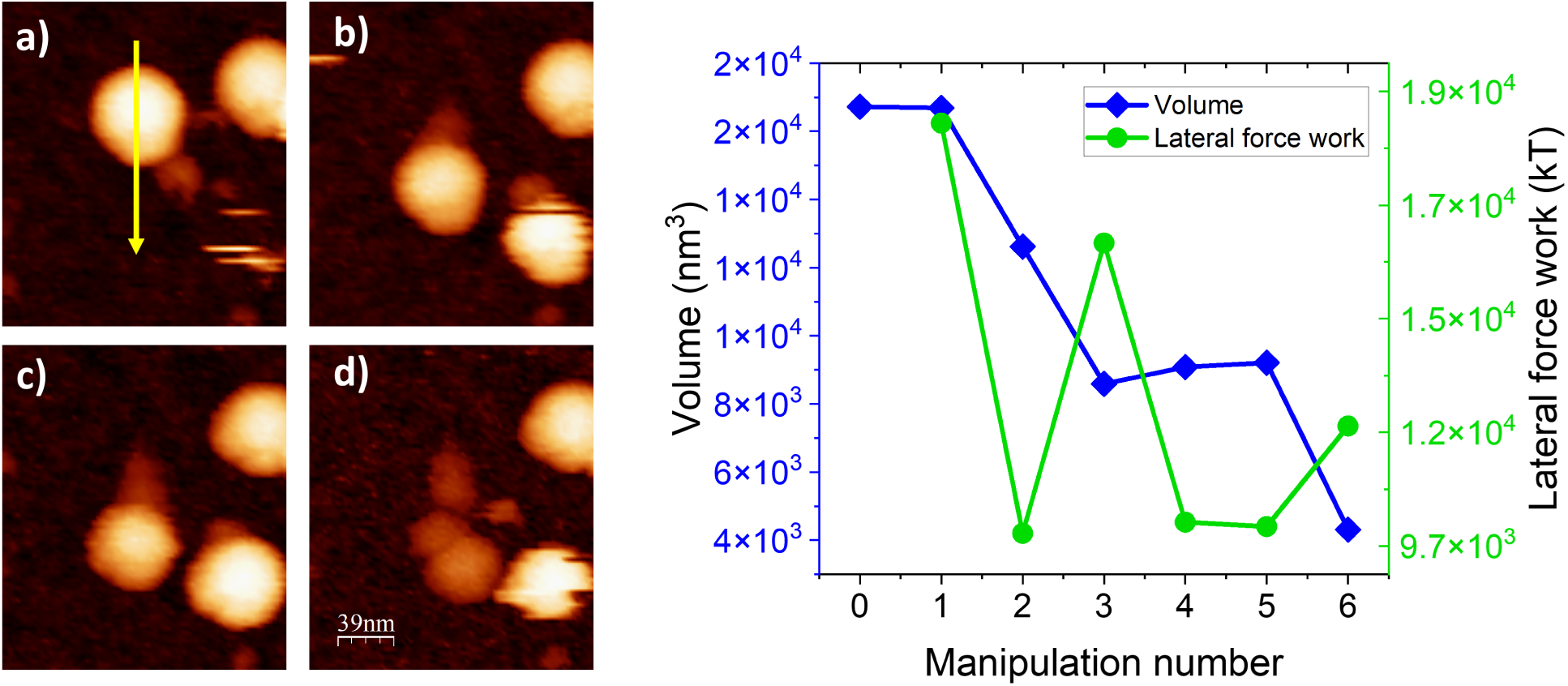
Topographic images of a BMV particle adhered to HOPG throughout a series of manipulations: (a). The particle at left begins fully intact and is pushed by the probe from top to bottom as indicated by the yellow arrow. (b,c) Repeated manipulations lead to a consistent decrease in particle height and (d) cause the formation of a trail of loose proteins until the particle is reduced to a small, loose pile of proteins. (e) Particle volume and lateral work force vs. manipulation number for a series of repeated manipulations performed on one BMV particle.

The trail of debris left by the particle after each manipulation (Fig. 10(a-d)) is a result of strong adhesive forces at the particle/substrate interface inducing capsomer extraction when the particle is moved. The resulting capsid lattice vacancies probably contribute to a weakening of the overall capsid structure, resulting in a cascading effect which eventually results in virion collapse/disassembly as seen in the final image of the series, Fig. 10(d).

The particle volume and lateral work force throughout the series of manipulations in (Fig. 10(a-d)) are shown in (Fig. 10(e)). From one scan to another, as the particle disassembles and its volume decreases, the lateral work force decreases as well. This indicates that intact particles are generally more resistant to lateral manipulation in comparison to severely structurally disrupted particles.

Changes in environmental conditions can impact the physicochemical properties underlying biological adhesion significantly. For example, the adhesion and hydrophobicity of algal cells are significantly affected by temperature and salinity^62–64^.

Viruses are sensitive and respond to chemical changes in their environment, too. Specifically, they can prevent or slow down decay when outside a cell host and yet promptly present their genome for replication after entry. Although this work’s primary aim is narrower than that of revealing the entire complexity of parameter landscape affecting static and dynamic virus adhesion, the methods outlined here could benefit investigations seeking to unveil virus response to environmental cues. As an example, we have asked how does the energy required to initiate the mechanical disassembly of BMV by rolling on HOPG vary with the pH of the solution? Fig. 11 shows that the magnitude of mechanical work to capsid disruption decreases almost an order of magnitude with increasing solution pH from 4.6 to 7.9. This can be expected because capsid cohesive interactions in cowpea chlorotic mottle virus (CCMV), a close relative of BMV, weaken as pH increases^65,66^.

**Figure 11:**
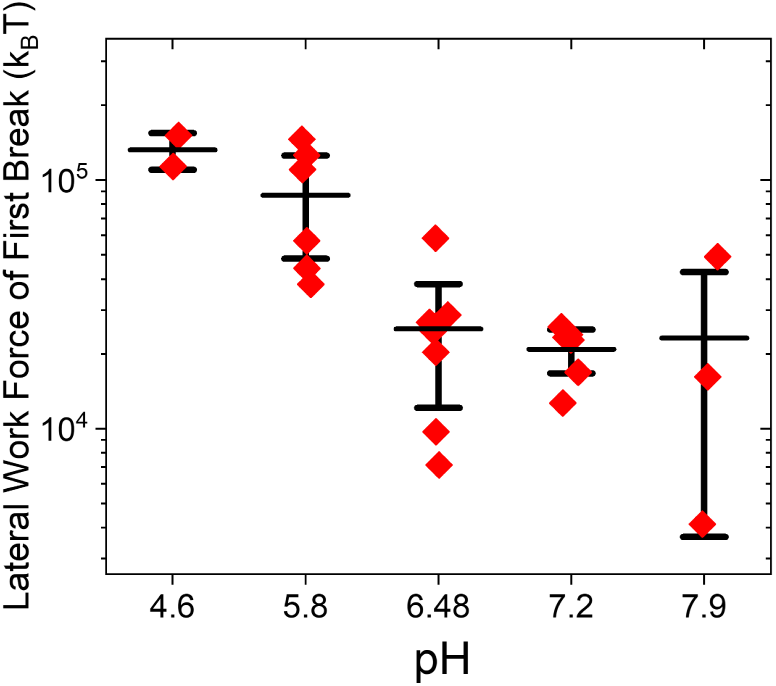
Lateral mechanical work leading to capsid disruption vs. pH in BMV on HOPG.

We now turn our attention to the dynamic response to lateral force manipulation of viruses vs. that of the rigid Au NP. In experiments with 80-nm AuNP in HOPG (Fig. 2), the lateral force rapidly reached ∼ 40 nN, which is 4–5 times higher than the average maximum lateral force measured in BMV, and 2–3 times the force measured on MPyV, Fig. 12. Therefore, although the lever arm of the lateral force torque was almost double for the 80 nm Au NP compared to the BMV, the resistance to displacement is greater in Au NPs. We note that the smaller, 24 nm Au NPs coated with CTAB, showed comparable if not slightly larger resistance to lateral force manipulation than the 80 nm Au NPs coated with PDADMAC (see Fig. S3). If the difference between the two AuNPs sizes was mainly due to contact area, the 80 nm AuNPs would have shown greater resistance to lateral force manipulation than the 24 AuNPs. Thus, we believe that the chemical nature of the ligand played a dominant role here, possibly through stronger hydrophobic interactions of the PDADMAC ligand with the HOPG substrate.

**Figure 12:**
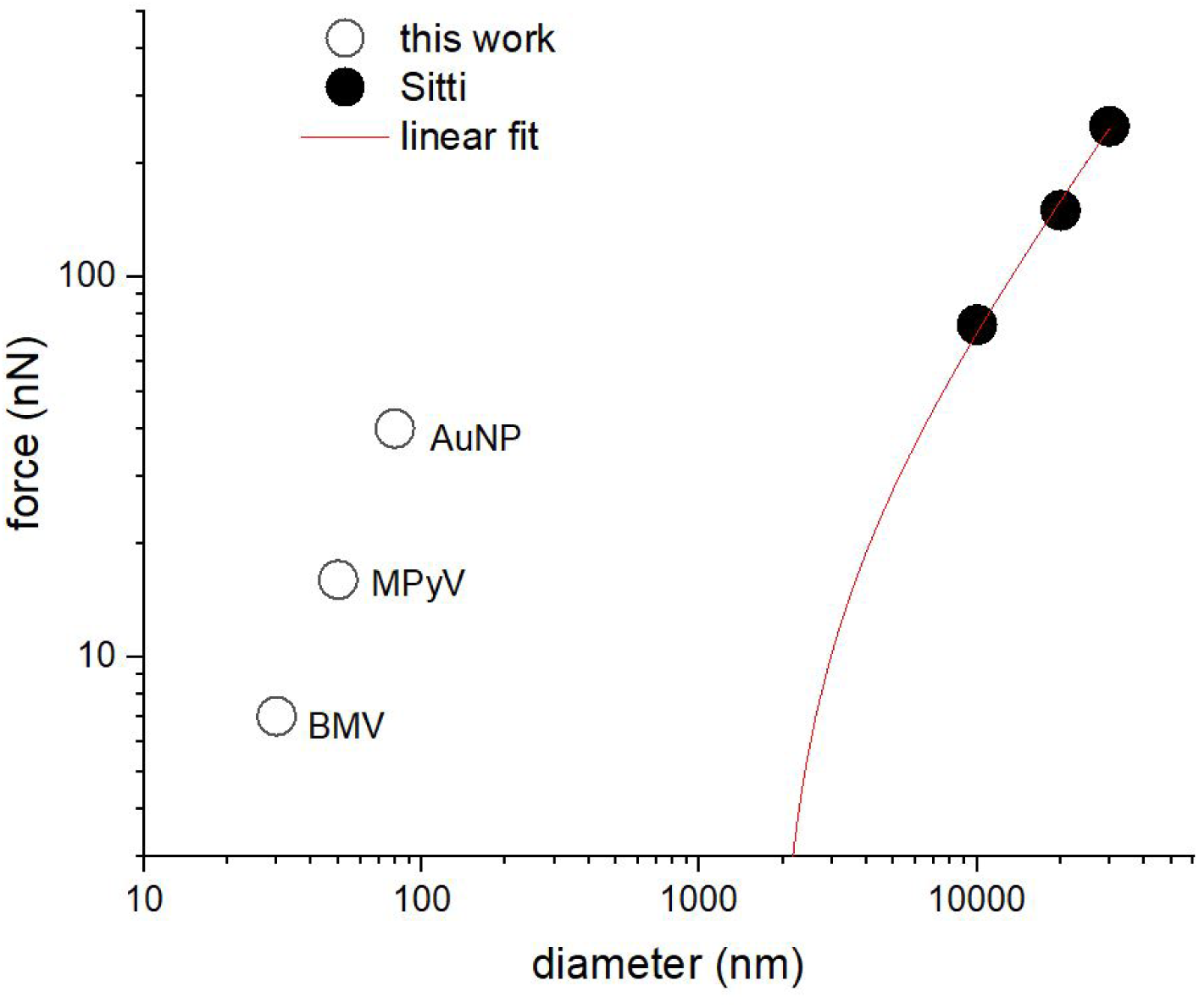
Quasi-static lateral force vs. nominal particle diameter for the three types of particles in this work and for previously published results from ref.^38^. The dash line (blue) represents a linear fit to the Sümer and Sitti data, extrapolated to the diameter range studied in this article.

For the weakly bound 80 nm Au particles, we observe the lateral force maximum occurring in the first half of the manipulation path (Fig. 2) while in virus particles, which are mechanically compliant, the maximum is generally observed in the second half of the manipulation path – this might be because virions are able to undergo greater deformation relative to particle size before enough force is applied to induce particle displacement.

It would be interesting to test whether the results obtained here in a liquid might follow a common scaling law with those obtained by Sümer and Sitti from AFM manipulation of polystyrene microparticles in air^38^. According to the Derjaguin theory for incompressible spherical particles of radius R in vapor on a flat surface^38^, the adhesion force scales linearly with the radius of the particle (eq. 5).

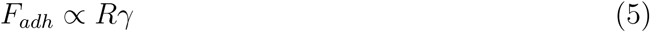

where *γ* stands for the interfacial energy (approx. half the work of adhesion^23^), typically few tens of mJ/m^2^. Assuming that friction force and adhesion force are also linearly proportional, we fit the force vs. diameter results by Sümer and Sitti from polystyrene microparticles in air^38^ with a linear fit and we extrapolate to the nanoscale for comparison with our results for viruses and Au NP in solution. Figure 12 shows the significant differences between, one hand – the lateral forces on viruses *vs.* nanoparticles of the same scale, and rigid nanoparticles in liquid *vs.* microparticles in air, on the other hand.

Incompressible Au NPs in water fall well above the extrapolation of data from Sümer and Sitti. Thus, AuNPs adhere more tenuously to the HOPG surface than one might expect, especially since it is well-known that water weakens the adhesion of solids. The opposite appears to happen here, possibly because of the hydrophobic effect which pushes the HOPG and AuNP together (AuNPs are surface functionalized with Au surface adsorbed polymer with polar groups for solubility, but the ligand coating is likely patchy).

Viruses cannot be considered incompressible at adhesion^3^. Upon contact, they deform elastically under the influence of attractive surface forces. The Johnson, Kendall, and Roberts (JKR) theory of contact mechanics^22^ provides a formal estimate for the radius of the area of contact between an elastic sphere and a flat surface:

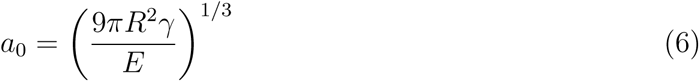

with the adhesion force estimated as:

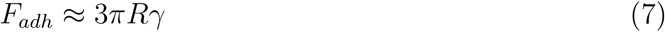

It is likely that friction is related to adhesion ^67^. If they are proportional, using JKR we can estimate both the ratios between Au NPs and viruses of the interfacial energy *γ* and the area of contact with the surface. To do so, we use (7) to obtain the coefficient between the interfacial energies for Au NPs and MPyV on HOPG like 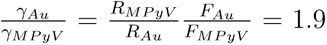, where *F_Au_* and *F_MP yV_* are the friction forces for Au NPs an MPyV particles, respectively. A similar estimation can be applied to find 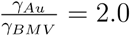. Thus, the interfacial energy of Au NPs is twice that of virus particles. Further, we can use the expression (6) to find the ratios of the contact radius with the surface between the compliant viruses and the stiff Au NPs. In the of MPyV particles we can derive 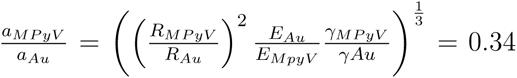 latter we used the Young Modulus for Au NPs (*E_Au_*) and MPyV virus (*E_MP yV_*) as 80 GPa and 0.5 GPa, respectively. For the case of BMV, this coefficient results in 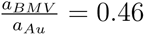. This means that the area of contact with the surface is 9 and 4 times larger for MPyV and BMV, respectively, than for the Au NPs. Thus, despite the fact that compliant viruses deform on the surface increasing their area of contact significantly with respect to the Au NPs, they remain less adhered than large and rigid particles. In this case, the higher interfacial energy of Au NPs takes over the geometrical facts of size and area of contact. Now it is pertinent to discuss the forces which are responsible of the interfacial energies of both viruses and Au NPs with the surface. Virus structures are populated with appendages, such as fibers and spikes, which are responsible for the interaction of viruses on the host cell surface. These interactions take place by specific bonds which depend on the biochemistry of the virus appendages and the host receptors ^68^. In our case surfaces are not functionalized with any receptors or antibodies and we only expect nonspecific interactions between the virus capsid and the surface. Consequently, we can assume to deal with non-specific physisorption governed by DLVO interactions as it happens with other biomolecules ^69^. DLVO interaction *F_DLV O_* include electrostatic *F_el_* and van der Waals forces *F_vdW_* as 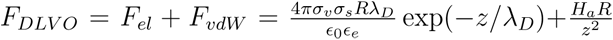. In this expression *σ_v_* and *σ_s_* are the charge densities of virus and surface, respectively; *ɛ*_0_ and *ɛ_e_* the electrical permitivities of vacuum and liquid, respecitvely; *z* the distance between the virus and the surface and *H_a_* the Hamacker constant. The Debye length *λ_D_* characterizes the exponential decrease of the potential resulting from screening the surface charges with electrolytes, with *λ_D_*= 0.174*/* [*C*] for a concentration *C* of divalent (1:2 or 2:1) electrolytes. In our case, the divalent concentration of the SAMA buffer is 0.008M ^3^, therefore *λ_D_* = 2*nm*. This means that the electrostatics is rapidly killed and the particles are attached to the surface through inespecific van der Waals forces. A similar discussion can be elaborated for Au NPs, indicating that van der Waals forces are more intense for Au NPs than for protein capsids.

The relative trends shown in Fig. 12 remained the same when mica replaced HOPG as a substrate. Specifically, MPyV started moving at 2-3 times the force required for BMV on both substrates. The adhesion of particles to both substrates is believed to be primarily the result of dispersion forces, while the affinity of particles to mica has an added contribution from long-range electrostatic forces ^69^. Virus-specific differences on the two substrates were relatively small: MPyV adhered somewhat (25%) stronger to HOPG than to mica, while for BMV there was practically no difference between mica and HOPG.

A molecular interpretation of differences between BMV at MPyV is difficult due to the fact that, in general, the virus surface is a mosaic of polar and non-polar residues as well as flexible and rigid protein domains, which all contribute to non-specific adhesion. We note the presence of large, flexible loops at the surface of MPyV^70^, which are normally involved in MPyV entry by receptor mediated endocytosis, while for BMV, which is a plant virus, such surface loops are lacking. BMV is believed to enter plant cells through external perforation damage of the cell walls.

An intriguing conclusion from these experiments is that, although mechanically compliant and therefore prone to realize a wider contact area, viruses show less resistance to rolling on any of the surfaces tested than rigid, ligand-stabilized Au NPs. This counterintuitive observation highlights the need for an in-depth theoretical treatment. At the same time, it raises the question: Might low resistance to rolling be a built-in capability aimed at increasing the rate of receptor binding and thus accelerating kinetics of cellular binding/entry? For now, this remains an open question which could receive an answer in the future by experiments similar to those presented here. It would be interesting to see how these data will change when known-specific ligand-receptor interactions are introduced on a model cell surface, such as a supported lipid bilayer. The preliminary work presented here indicate that such explorations should be possible.

## Conclusion

Lateral force manipulation by AFM was explored as a possible approach for comparative studies of adhesion interactions in isometric viruses in particular and other mechanically-compliant nanoparticles in general. Virus-probe and virus-support interactions tend to be complex; as such, multi-pronged approaches that address different properties should be utilized. With its set of complementary, simultaneous read-out variables, liquid-cell AFM is a suitable method. It was found that, under physiological buffer conditions, non-enveloped virions of an animal and a plant virus adhere non-specifically in such a way that they can be manipulated on the surface over extended distances relative to their radius without desorption. The finding will be valuable in designing new experiments seeking to understand virus mobility on biological surfaces.

MPyV and BMV exhibited distinct responses from rigid Au NPs to lateral force manipulation. Although MPyV is several times more resistant to rolling than BMV, the resistance to rolling in both viruses is significantly lower than that of rigid Au NPs, despite the fact that viruses, being mechanically compliant, presumably have a contact area more extended than that of rigid nanoparticles of similar diameter. The mechanical work lateral force measured during the rolling motion of viruses is an order of magnitude greater than the estimated free energy of the assembly of capsids, suggesting a high barrier against disassembly by mechanical forces. Significant virus particle strain was observed during lateral force manipulation for both viruses and should be considered in future modeling seeking to extract information about virus interfacial interactions during the virus life-cycle.

## Supporting Information Available

Contains information on capsid structures for BMV and MPyV.

## Conflicts of Interest

None.

## Data Availability

Data freely available by direct request to corresponding authors or via publicly accessible IU Scholar Works Repository: https://libraries.indiana.edu/databases/scholarworks.

## Acknowledgement

The work was partially supported by the U.S. Army Research Office through award #W911NF1310490 and #W911NF-24-1-0225 to B.D.. B.D. acknowledges support from project grant # PNRR-I8/C9-CF105 under contract # 760099 from the Romanian Ministry of Research, Innovation, and Digitization. P. J. de Pablo acknowledges the Salvador de Madariaga Scholarship PRX21/00317.

The authors thank Dr. Xingchen Ye for the CTAB-coated Au nanoparticle samples. C.A. also thanks the Indiana Grad Workers Coalition – United Electric Workers for its support and representation.

## Notes

### Competing Interest Statement

The authors have declared no competing interest.

