## Supplemental Information for "Nanotribology of Viruses Reveals Their Adhesion Strength and Modality of Motion on Surfaces"

### Capsid structures

MPyV and BMV have both icosahedral protein shells (capsids) protecting their genomes. The two capsids may share the same overall symmetry, but they are different in the type of cargo they encapsulate (single stranded RNA for BMV, double stranded DNA for MPyV), and the number and nature of capsomers. Their capsid structures are presented, at the same scale, in Fig. S1. Structures were constructed from ViperDB v3.0[1] data sets constructed from PDB coordinates of the VP1 MPyV protein[2] and the BMV coat protein (PDB structure 3J7L)[3].

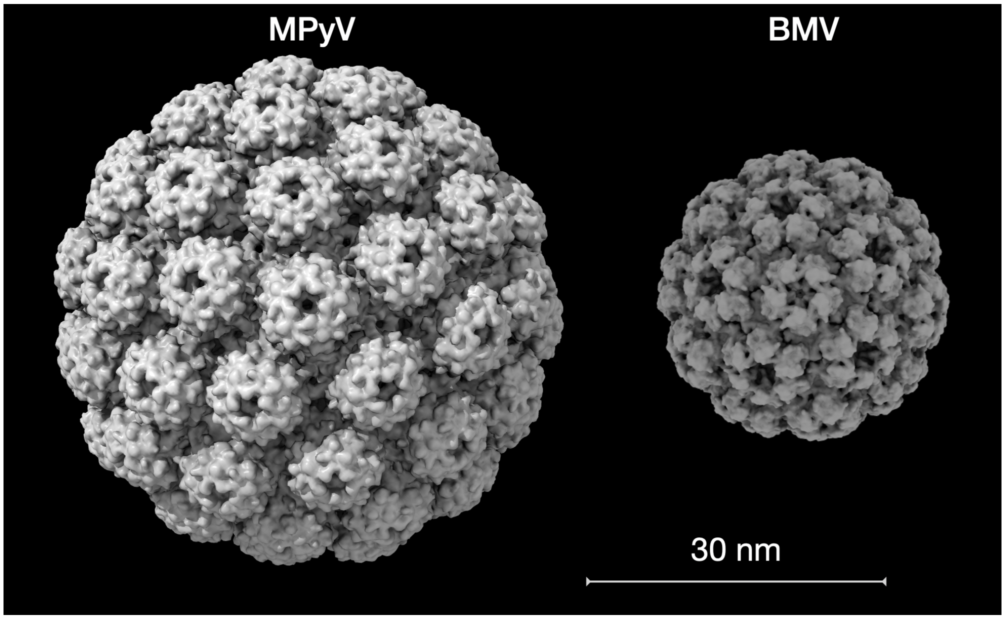


Figure S1: Structural models of murine polyoma and brome mosaic virus capsids.

Raw data for lateral signal during particle manipulation are presented in Figs. S3, S4 and S5. For data processing particle coming in contact with tip is found when amplitude drops at least 2 times and set as 0 distance, then average background before contact is subtracted for all traces. All traces shown were collected from a single experiment, on the same sample preparation with the same tip, to provide experimental variations not affected by sample preparation or different tip spring constant.

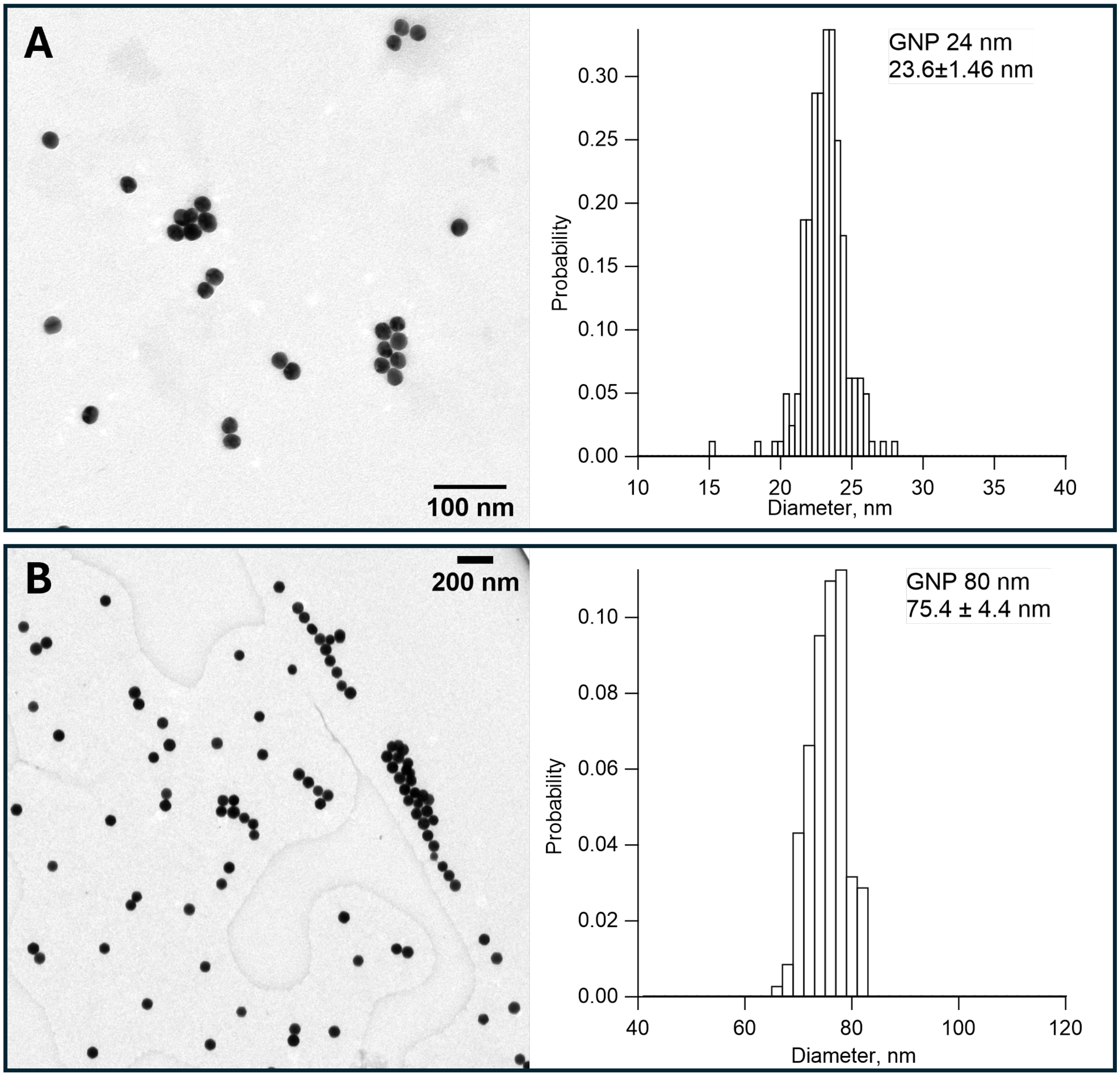


Figure S2: TEM images and size histograms of GNP 24 nm (A) and 80 nm (B) samples. 10 uL of sample was placed on carbon coated TEM grid and excess of solution was blotted by filter paper after 5 min. Dried samples were imaged on JEOL 1010.

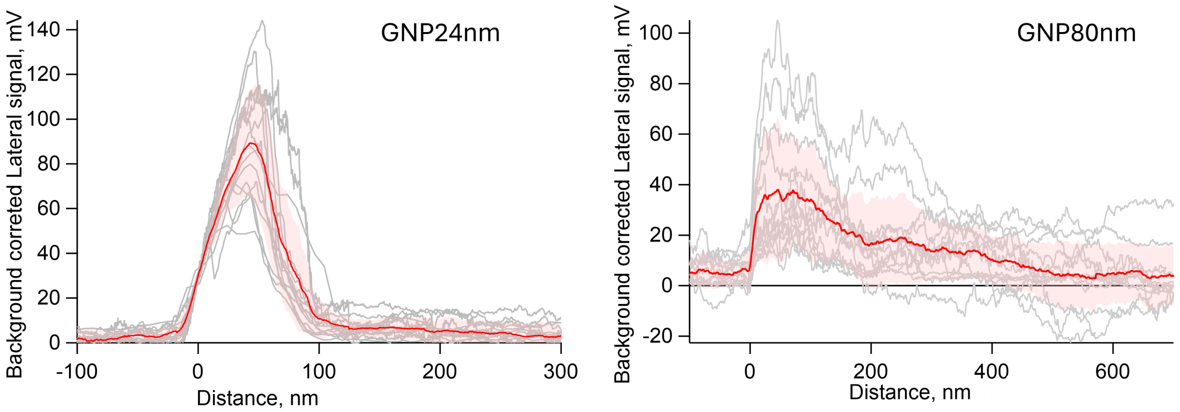


Figure S3: Raw data for lateral signal from gold nanoparticles (GNPs) on HOPG having diameters of 24nm and 80 nm. The two GNPs have different types of ligand coats (see Experimental section, main text), with the GNP24nm coat being more hydrophobic, and being expected to bind more tenuously to HOPG. Gray traces: individual measurements. Pink background: standard deviation. Red: average trace.

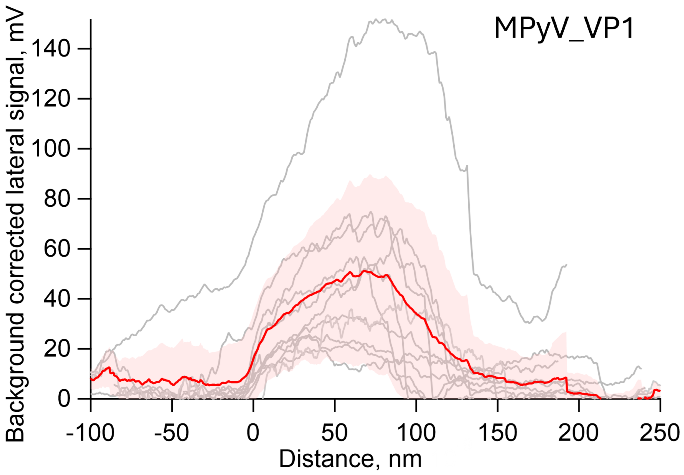


Figure S4: Raw data for lateral signal for MyPV VLPs on HOPG. All traces obtained with the same tip, in the same day, and buffer.

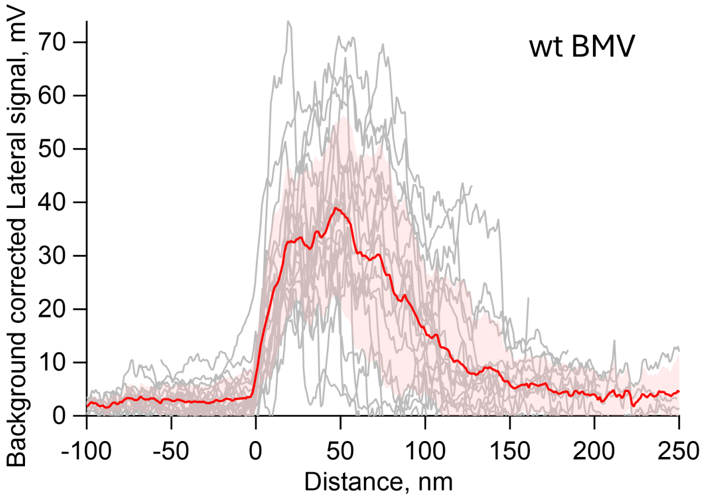


Figure S5: Raw data for lateral signal for wtBMV.

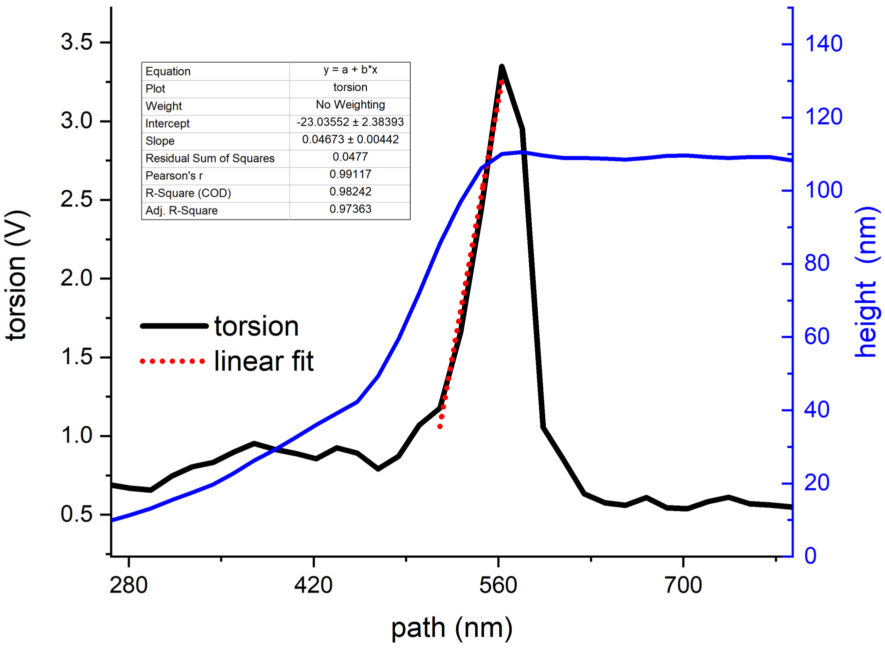


Figure S6: Calibration of lateral force scanning the tip on the step of a grid, keeping the cantilever parallel to the step. The linear fit of the lateral torsion through the step results in $0.047V/nm$ $(21.3nm/V)$. For example, in order to obtain the maximum lateral force of the red curve in Fig. S3, this calibration is multiplied by $0.05V$, resulting in a lateral bending of $1.2nm$. In turn, using formula (1) $k_l = 5.3N/m$, about $\sim 60$ times larger than $k_n$. Therefore, $F_l = 5.3N/m \times 1.2nm = 5.6nN$.
